# A force-sensitive adhesion GPCR is indispensable in normal equilibrioception

**DOI:** 10.1101/2023.10.24.563886

**Authors:** Zhao Yang, Shu-Hua Zhou, Qi-Yue Zhang, Zhi-Chen Song, Wen-Wen Liu, Yu Sun, Ming-Wei Wang, Xiao-Long Fu, Kong-Kai Zhu, Ying Guan, Jie-Yu Qi, Xiao-Hui Wang, Yu-Nan Sun, Yan Lu, Yu-Qi Ping, Yue-Tong Xi, Zhi-Gang Xu, Wei Xiong, Wei Qin, Wei Yang, Fan Yi, Xiao Yu, Ren-Jie Chai, Jin-Peng Sun

**Affiliations:** NHC Key Laboratory of Otorhinolaryngology, Qilu Hospital of Shandong University, and New Cornerstone Science Laboratory, Department of Biochemistry and Molecular Biology, School of Basic Medical Sciences, Cheeloo College of Medicine, Shandong University, Jinan 250012, China; State Key Laboratory of Digital Medical Engineering, Department of Otolaryngology Head and Neck Surgery, Zhongda Hospital, School of Life Sciences and Technology, Advanced Institute for Life and Health Jiangsu Province High-Tech Key Laboratory for Bio-Medical Research, Southeast University, Nanjing 210096, China; Key Laboratory of Experimental Teratology of the Ministry of Education, Department of Physiology, School of Basic Medical Sciences, Cheeloo College of Medicine, Shandong University, Jinan 250012, China; Department of Otolaryngology-Head and Neck Surgery, Shandong Provincial ENT Hospital, Cheeloo College of Medicine, Shandong University, Jinan, 250014, China; Department of Otorhinolaryngology, Union Hospital, Tongji Medical College, Huazhong University of Science and Technology, Wuhan, China; Advanced Medical Research Institute, Cheeloo College of Medicine, Shandong University, Jinan 250012, China; Department of Clinical Laboratory, The Second Hospital, Cheeloo College of Medicine, Shandong University, Jinan 250012, China; Medical Science and Technology Innovation Center, Shandong First Medical University & Shandong Academy of Medical Sciences, Jinan 250000, China; Key Laboratory of Molecular Cardiovascular Science of the Ministry of Education, Department of Physiology and Pathophysiology, School of Basic Medical Sciences, Peking University, Beijing 100191, China; Shandong Provincial Key Laboratory of Animal Cells and Developmental Biology, Shandong University School of Life Sciences, Qingdao 266237, China; Chinese Institute for Brain Research, Beijing 102206, China; School of Physics, State Key Laboratory of Crystal Materials, Shandong University 250012, Jinan, China; Department of Biophysics, and Department of Neurology of the Fourth Affiliated Hospital, Zhejiang University School of Medicine, Hangzhou 310058, China; Department of Pharmacology, School of Basic Medical Sciences, Shandong University, Jinan, 250012, China; Co-Innovation Center of Neuroregeneration, Nantong University, Nantong 226001, China

## Abstract

Equilibrioception is essential for the perception and navigation of mammals in the three-dimensional world. A rapid mechanoelectrical transduction (MET) response in vestibular hair cells plays a critical role in positional and motional perception. Here, we identified that the G protein-coupled receptor LPHN2/ADGRL2, which is expressed in the apical membrane of utricular hair cells, is required for maintenance of normal balance. Hair cell-specific Lphn2 deficiency in mice impaired both balance behaviors and MET responses. Functional analyses using *Pou4f3-CreER^+/-^*; *Lphn2^fl/fl^* mice and LPHN2-specific inhibitors revealed that LPHN2 regulated the tip link-independent MET current at the apical surface of the utricular hair cell by converting force stimuli into transmembrane channel-like protein 1 (TMC1) activity. Force sensation by LPHN2 also induced glutamate release and calcium signaling in utricular hair cells. Reintroduction of LPHN2 into the hair cells of Lphn2-deficient mice restored the vestibular functions and MET responses. Our data suggest an indispensable role for a mechanosensitive GPCR in equilibrioception.

## Introduction

A sense of balance and motion enables our perception and navigation in the three-dimensional world and is thus critical to our interactions with the world. Positional or motional information is primarily perceived by hair cells located in the sensory epithelium of vestibular end organs, which include two perpendicularly arranged otolith organs, the utricle and saccule, and three semicircular canals^1,2^. Extensive evidence has shown that vestibular hair cells (VHCs) transmit balance information about head motion or tilt into electrical signals through hair bundle-deflection-gated ion channels opening, which is termed mechanoelectrical transduction (MET)^3,4^. Extremely rapid channel gating on a microsecond timescale occurred after hair bundle displacement in response to external forces, most likely through tip links, as illustrated by the seminal work of the Hudspeth laboratory on frogs and many others on other systems^3–6^. Most recently, two candidate channels, TMC1 and TMC2, which are expressed at the tip of stereocilia, have been suggested to be the pore-forming subunits of MET channels and were shown to form a complex with additional subunits, such as TMIE, LHFPL5, CIB2/3 and tip-link proteins, to sense and transduce mechanical forces^7–10^.

To date, the force sensation in the MET has been hypothesized to be mediated by tip link components or TMC1/2^5,11,12^. In addition to ion channels, membrane receptors belonging to the family of G protein-coupled receptors (GPCRs), which are the most common drug targets, can sense mechanical forces^13–19^. For example, several adhesion GPCRs (aGPCRs), which possess large and multidomain N-termini that allow receptors to interact with extracellular matrices, can respond to mechanical force stimulation which have been shown by our labs and several others ^18,20–24^ (Supplementary information, Fig. S1a). Ion channels and GPCRs are two classes of sensory receptors. Notably, temperature and touch are sensed by TRP channels and Piezo channels, respectively^25,26^. In contrast, vison and olfaction are mainly mediated by GPCRs, which convert light or odor stimuli into electrical signals through coupling to cyclic nucleotide-gated (CNG) channels^27–30^. Both GPCR family members and ion channels participate in distinct taste sensations^31,32^. The downstream signaling and cellular outputs differ between GPCRs and ion channels. Whereas sensory ion channels directly mediate ion permeability, GPCRs regulate sensory signals by controlling the intracellular concentrations of secondary messengers (cAMP, cGMP, Ca^2+^, etc.). The generation of a second messenger requires enzyme catalysis, the speed of which is normally limited by diffusion limits. Therefore, despite playing important roles in the sensation of light, smell and taste, the GPCR-second messenger system is conventionally excluded from the equilibrioception process in hair cells due to its relatively slow kinetics (tens to hundreds of milliseconds).

Although the ion-channel-centered MET system may play central roles in equilibrioception, it may not depict the full landscape of our balance perception. Importantly, many types of movement generally require hundreds of milliseconds to seconds to reach equilibration, such as walking, dancing, running, accelerating a car to 100 miles/hour or diving. Therefore, equilibrioception may not only be detected very rapidly but may also be perceived over a longer period. Moreover, GPCRs may regulate the activity of ion channels by direct physical interactions and conformational transitions, thus bypassing the time-consuming second messenger system. We thus cannot exclude the possibility that, in addition to ion channels, GPCRs may also actively participate and play modulatory roles in equilibrioception (Fig. 1a). We speculated that these equilibrioception receptors should fulfill the following criteria: (1) they are expressed in the stereocilia or apical membrane of VHCs; **(2)** they can directly sense force in a physiological range (2∼100 dynes/cm^2^); **(3)** they are able to convert force stimuli into chemical or electrical signals in VHCs or neurotransmitter release from VHCs; and **(4)** genetic ablation of these receptors in animal models leads to equilibration disorders.

**Fig. 1.**
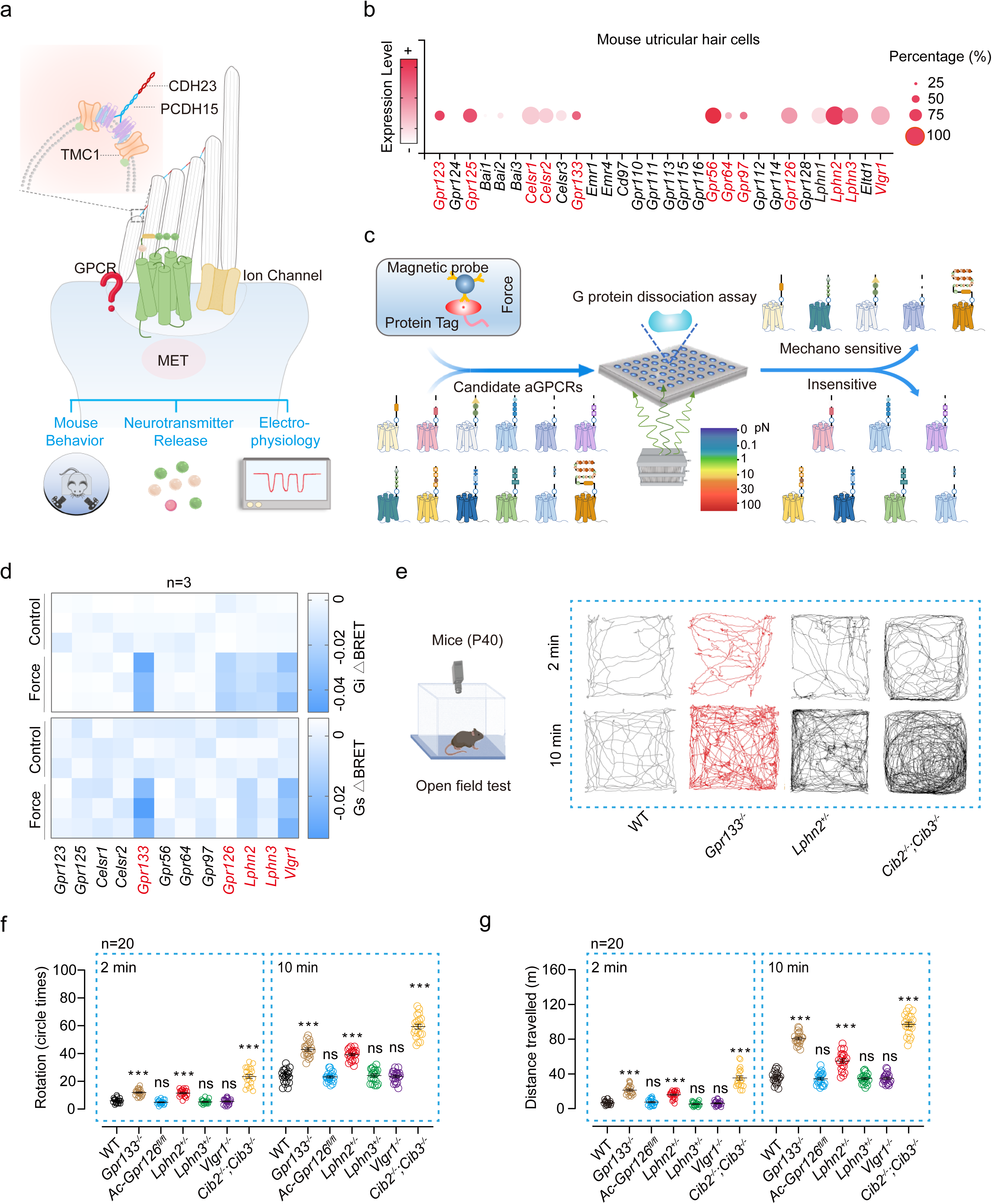
Screening of mechanosensitive adhesion GPCRs in vestibular hair cells **(a)** Schematic representation of the potential roles of ion channels and GPCRs in MET process in vestibular hair cells. Enlarged box shows the tip link, composed of PCDH15 and CDH23, and putative components of the MET channel complex at the tip of stereocilia. **(b)** Expression profiles of 30 adhesion GPCR (aGPCR) genes in mouse utricular hair cells. Single cell RNA-seq datasets of utricular hair cells are from GSE71982. The size of each circle represents the percentage of hair cells where the gene is detected, and the color intensity represents the average mRNA expression level of the gene. The 12 aGPCRs expressed in more than 20% of the utricular hair cells are highlighted. **(c)** Schematic representation of the strategy used to screen the mechanosensitive aGPCRs. **(d)** Summary of the force-induced Gi3 (top panel) and Gs (bottom panel) activation of 12 aGPCRs. A force of 10 pN was applied to the receptors and the Gi3 or Gs activation was measured by BRET assay, which was presented as a heatmap (n=3). **(e)** Schematic view (left) and representative tracks (right) of WT, *Gpr133^-/-^*, *Lphn2^+/-^* and *Cib2^-/-^; Cib3^-/-^* mice in open-field test during 2 min or 10 min tracking period. **(f, g)** Quantification of the circling (left) and travelling activity (right) of WT, *Gpr133^-/-^*, *Gpr133^+/-^*, *Atoh1*-*cre*^+/-^*Gpr126^fl/fl^* (referred to as *Ac*-*Gpr126^flfl^*), *Lphn2^+/-^*, *Lphn3^+/-^*, *Vlgr1^-/-^* and *Cib2^-/-^; Cib3^-/-^*mice in open-field test (n=20 mice per group). (**f, j**) ***P < 0.001; ns, no significant difference. Gene knockout mice compared with WT mice. The bars indicate mean ± SEM values. All data were statistically analyzed using one-way ANOVA with Dunnett’s post hoc test.

According to the above criteria for equilibrioception receptors, in the present study, we screened the mechanosensitivity of adhesion GPCRs (GPCRs with force-sensing potential) in utricular hair cells via expression level analyses, cellular assays and animal models. We identified two GPCRs, LPHN2 and GPR133, as mechanosensitive receptors in the vestibular system that are indispensable for equilibrioception. Specifically, LPHN2 was expressed at the apical surface of over 80% of utricular hair cells, and global knockout (KO) or inducible hair cell-specific KO of *Lphn2* impaired balance behaviors in mice. Moreover, the MET current in utricular hair cells was impaired either by genetic ablation of *Lphn2* or by pharmacological inhibition of LPHN2 with a specific inhibitor in a reversible manner. Mechanistically, LPHN2 coupled to TMC1 expressed at the apical surface of utricular hair cells and specifically participated in the normal-polarity MET, which is independent of that mediated by the tip link of stereocilia. Furthermore, force sensation by LPHN2 in utricular hair cells induced a Ca^2+^ response and glutamate release, the magnitude of which is approximately half of that induced by force applied on tip link component CDH23. Collectively, our findings suggest that a mechanosensitive GPCR is required for normal balance; this receptor actively regulates a previously uncharacterized MET process at the apical surface of VHCs by functionally coupling to TMC1.

## Results

### Screening of mechanosensitive adhesion GPCRs in the vestibular system

The human aGPCR family consists of 33 members (30 in mice), and several members have been reported to be activated by mechanical forces (Supplementary information, Fig. S1a)^17,18,21,24,33–35^. Single-cell RNA sequencing (scRNA-seq) data indicated that 12 aGPCRs were expressed in more than 20% of the utricular hair cells of the mice (Fig. 1b). We then established a high-throughput mechanical stimulation assay to examine the mechanosensitive potential of these 12 aGPCRs that were enriched in the vestibular system using a magnetic tweezer system connected to a GPCR biosensor platform. Briefly, tensile forces were applied to paramagnetic beads coated with anti-Flag M2 antibody or polylysine using a magnetic system, and force-induced G protein activation was examined using a bioluminescence resonance energy transfer (BRET) assay in HEK293 cells transfected with plasmids encoding N-terminal Flag-tagged GPCR and G protein biosensors (Fig. 1c)^36,37^. In response to magnetic stimulation, tension force is produced by magnetic beads attached to the N-terminus of selected aGPCRs, and the activation of Gs or Gi signaling downstream of mechanosensitive GPCRs is measured by BRET signals, which are well established methods for detecting GPCR activation^18,20,37–41^. Specifically, a force of 10 pN was applied via magnetic beads, which equaled 100 dynes/cm^2^ on the plasma membrane and was adopted in a previous study to mimic arterial wall shear stress (approximately 1 m/s under physiological conditions and similar to the normal walking pace)^42^. Using this system, we revealed that 5 receptors (GPR133, GPR126, LPHN2, LPHN3 and VLGR1) activated Gi signaling in response to force stimulation, whereas 3 of them (GPR133, LPHN2 and VLGR1) also activated Gs signaling in response to force application (Fig. 1d).

We then investigated whether these mechanosensitive aGPCRs are indispensable for equilibrioception by comparing *Gpr133*^-/-^ mice, *Lphn2*^+/-^ mice, *Lphn3*^+/-^ mice, *Atoh1*-*Cre*^+/-^;*Gpr126^fl/fl^*mice and *Vlgr1*^-/-^ (*Vlgr1*/del7TM) mice with wild-type (WT) littermates in the open field test, which is a commonly used assay for evaluating vestibular functions (Fig. 1e-g). *Lphn2*^+/-^ mice and *Lphn3*^+/-^ mice were used since homozygous KO of *Lphn2* or *Lphn3* caused embryonic lethality and developmental defects, respectively^43,44^. *Gpr126* deficiency also leads to embryonic lethality due to cardiac abnormalities; therefore, we generated *Atoh1*-*Cre*^+/-^;*Gpr126^fl/fl^* mice to eliminate Gpr126 expression in hair cells (*Atoh1* is a marker gene for nascent cochlear and VHCs^45,46^, and scRNA-seq data suggested that approximately 85% of Gpr126-expressing utricle hair cells have detectable *Atoh1*). The deficiency of the target receptor genes in the mutant mice was verified by genotyping PCR and western blotting analysis (Supplementary information, Fig. S1b-i; Fig. S2a-j). All the above mice were viable, fertile (except for the *Gpr133^-/-^* female mice that were sterile) and had normal body weights under normal chow diet conditions (Supplementary information, Fig. S2k, l). Notably, with *Cib2*^-/-^;*Cib3*^-/-^ double mutant mice as a positive control, behavioral analyses of these mice in the open field test revealed that the *Gpr133^-/-^* mice and *Lphn2*^+/-^ mice exhibited approximately 1.5-2-fold increases in circling behavior and traveling distances compared with those of their WT littermates in both the 2 min time frame (*Gpr133^-/-^*: 12.0 ± 0.6 circles and 21.4 ± 1.3 meters; *Lphn2^+/-^*: 11.6 ± 0.6 circles and 16.0 ± 1.0 meters; WT: 6.0 ± 0.4 circles and 7.0 ± 0.6 meters) and the 10 min time frame (*Gpr133^-/-^*: 42.9 ± 1.2 circles and 80.8 ± 1.6 meters; *Lphn2^+/-^*: 39.3 ± 0.9 circles and 54.9 ± 2.4 meters; WT: 23.8 ± 1.1 circles and 36.0 ± 1.5 meters). In contrast, the *Atoh1*-*Cre^+/-^*;*Gpr126^fl/fl^* mice, *Vlgr1*^-/-^ mice, and *Lphn3*^+/-^ mice did not show significantly abnormal circling or traveling behaviors compared with the WT controls (Fig. 1e-g). These results suggest that mechanosensitive LPHN2 and GPR133 may play a role in maintenance of normal balance.

### Mechanosensitive LPHN2 is indispensable for normal vestibular functions

Because a functional and mechanistic analysis of GPR133 was described in another study^47^, we focused on LPHN2 in the current manuscript. We next assessed the expression of LPHN2 in the vestibular system by quantitative RTLPCR (qPCR) analysis and revealed that LPHN2 exhibited constant and stable expression from late embryos (E15) to adulthood (P120) (Supplementary information, Fig. S3a). Both the scRNA-seq data and the results from our single-hair-cell qRTLPCR analysis confirmed the expression of LPHN2 in the utricular hair cells. Compared with LPHN3, another mechanosensitive LPHN family member expressed in utricular hair cells, LPHN2 exhibited both significantly greater apparent expression frequency and average expression level (Supplementary information, Fig. S3b-f).

The vestibular behaviors of the *Lphn2*^+/-^ mice at P40 were then assessed through the rotarod test and forced swimming test, with the *Cib2*^-/-^; *Cib3*^-/-^ double mutant mice as a positive loss-of-function control. Despite their nearly normal performance in the rotarod test (*Lphn2*^+/-^ 109.8 ± 3.7 s vs. WT 110.6 ± 4.0 s), the *Lphn2*^+/-^ mice exhibited impaired swimming ability, with a swimming score of 0.88 ± 0.13 (the 0–3 scoring system was used; WT mice and *Cib2*^-/-^; *Cib3*^-/-^ mice scored 0 and 2.71 ± 0.13, respectively) (Supplementary information, Fig. S4a, b). To specifically investigate the vestibular functions of the *Lphn2*^+/-^ mice, we measured their vestibular-ocular reflex (VOR) responses to sinusoidal head rotations at P40. Notably, compared with the WT mice, the *Lphn2*^+/-^ mice showed an approximately 25%-50% decrease in compensatory VOR gains in response to either earth-vertical or off-vertical axis rotation, suggesting a deficiency in both the semicircular canals and otolith system (Supplementary information, Fig. S4c-f). In contrast to the *Lphn2*^+/-^ mice, the *Lphn3*^+/-^ mice did not significantly differ from WT mice in the rotarod test, forced swimming test or VOR test; thus, *Lphn3*^+/-^ mice served as a negative control for vestibular behavior analyses (Supplementary information, Fig. S4a-f). Collectively, these results indicated that LPHN2 but not LPHN3 plays an important role in balance maintenance.

### Expression pattern of LPHN2 in utricular hair cells

We next examined the expression patterns of LPHN2 in the mouse utricle by whole-mount immunostaining and found that LPHN2 was specifically expressed in approximately 80% of the Myo7a-positive hair cells (Fig. 2a). The specificity of the LPHN2 antibody was supported by both the western blotting results in HEK293 cells expressing different LPHNs and the immunostaining of utricles derived from *Lphn2*^-/-^ mice at E18 (*Lphn2*^-/-^ mice showed embryonic lethality) (Fig. 2a; Supplementary information, Fig. S5a). The expression pattern of LPHN2 was further supported by RNAscope in situ hybridization, which revealed a comparable percentage of LPHN2-expressing hair cells (∼80%) to that determined by immunostaining (Fig. 2b, c). To further determine the subcellular localization of LPHN2 in utricular hair cells, we examined the expression pattern of LPHN2 in different optical sections spanning from the stereocilia to the hair cell body. LPHN2 expression was found only at the apical surface of the utricular hair cells, as revealed by its close proximity to spectrin, which is a specific marker of the cuticular plate in hair cells^48,49^ (Fig. 2d; Supplementary information, Fig. S5b). In contrast, LPHN2 expression was not found within the stereocilia or along the basolateral membrane of hair cells (Fig. 2d; Supplementary information, Fig. S5b). The specific expression of LPHN2 at the apical surface was further supported by section staining of the utricle sensory epithelium using Myo7a and Sox2 as the hair cell and supporting cell markers, respectively (Supplementary information, Fig. S5d). Consistent with the antibody-based expression analysis, we observed LPHN2-mCherry expression only at the apical surface of the utricular hair cells, utilizing an LPHN2^mCherry^ transgenic knock-in mouse line (Fig. 2e). The expression pattern of mechanosensitive LPHN2 at the apical membrane surface of utricular hair cells suggested that it might participate in force sensation during equilibrioception.

**Fig. 2.**
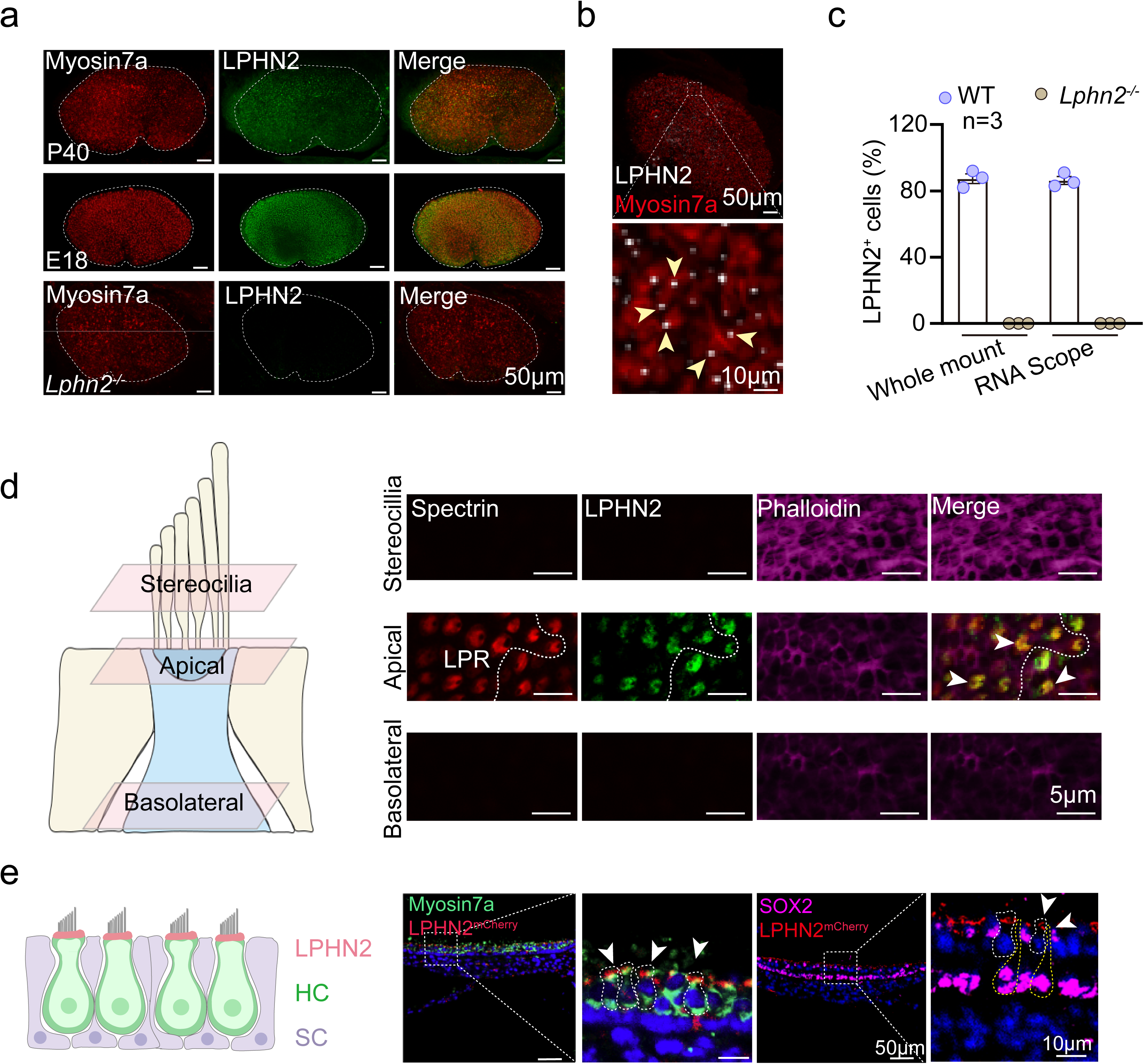
LPHN2 is specifically expressed at the apical surface of utricular hair cells **(a)** Coimmunostaining of LPHN2 (green) with Myosin7a (red) in utricle whole mounts derived from WT mice at E18 or P40, or from *Lphn2^-/-^* mouse embryos at E18 (n = 3 mice per group). Scale bar, 50 μm. **(b)** Representative images of whole-mount RNA scope *in situ* hybridization of LPHN2 (white) combined with immunostaining of Myosin7a (red) in the utricle of P40 mice (n = 3 mice). Arrows indicate LPHN2 staining in myosin7a-expressing hair cells. Scale bars, 50 μm and 10 μm for low- and high-magnification views, respectively. **(c)** Quantitative analysis of LPHN2 expression in myosin7a-positive utricular hair cells from WT mice or *Lphn2^-/-^* embryos. Data are correlated to Fig. 2a, b (n = 3 mice per group). **(d)** Left panel: Diagram of utricular hair cells showing the selected optical planes (stereocilia, apical surface or basolateral section) for imaging by confocal microscopy. Right panel: Coimmunostaining of spectrin (red), LPHN2 (green) and phalloidin (magenta) at different optical planes of hair cells in utricle whole mounts of P40 mice. Arrows indicate coimmunostaining of LPHN2 with spectrin. The line of polarity reversal (LPR) is depicted as a white dotted line. Scale bar, 5 μm. Data are correlated to Supplementary information, Fig. S5b, c. **(e)** Left panel: Schematic view of LPHN2 (red) expression in utricular hair cells. HC, hair cells; SC, supporting cells. Right panel: Expression of LPHN2-mCherry (red) with Myosin7a (green) or with SOX2 (magenta) in utricular sections derived from LPHN2^mCherry^ mice at P40 (n = 3 mice per group). Arrows indicate the distribution of LPHN2-mCherry at the apical surface of utricular hair cells. The utricular hair cells and supporting cell are depicted by while and yellow dashed lines, respectively. Scale bars, 50 μm and 10 μm for low- and high-magnification views, respectively.

### LPHN2 in VHCs specifically regulates balance sensation

To determine the specific functional role of LPHN2 in VHCs, we crossed *Lphn2^fl/fl^* mice with inducible *Pou4f3-CreER^+/-^* transgenic mice (Fig. 3a). Pou4f3 is the transcriptional target of ATOH1 and a commonly used marker of hair cells and is expressed in all detected LPHN2-positive utricular hair cells, as revealed by scRNA-seq data^50,51^ (Supplementary information, Fig. S5e, f). The specificity of LPHN2 ablation in the vestibule of the *Pou4f3-CreER^+/-^*; *Lphn2^fl/fl^* mice was indicated by the loss of LPHN2 immunostaining in the Pou4f3-expressing utricular hair cells but not in other tissues, such as the vestibular or somatosensory nuclei in the brainstem (Fig. 3b; Supplementary information, Fig. S5g-h). The specific decrease in Lphn2 expression in the vestibules of the *Pou4f3-CreER^+/-^*; *Lphn2^fl/fl^* mice was further supported by western blot analysis (Supplementary information, Fig. S5i).

**Fig. 3.**
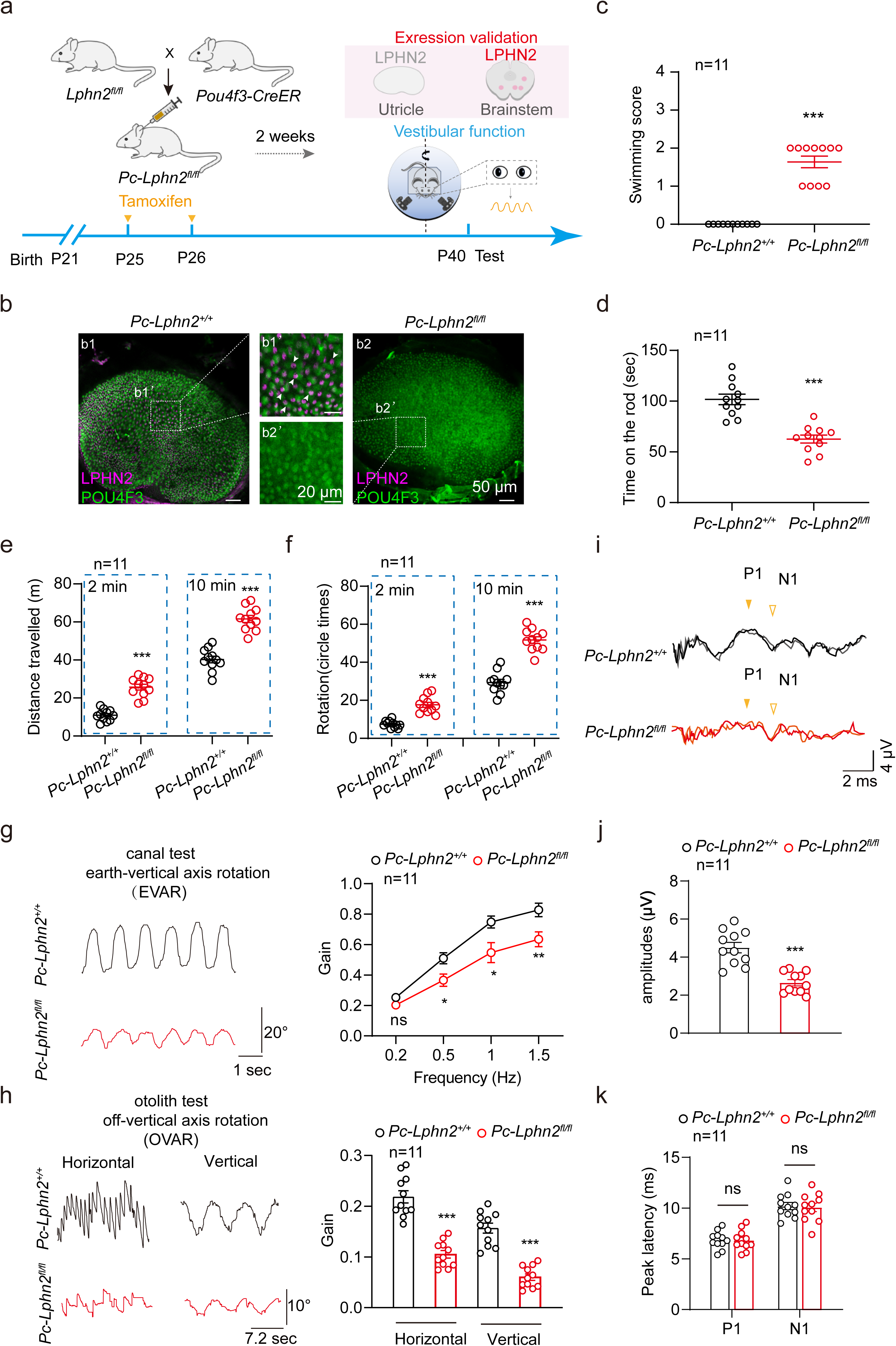
Conditional knockout of LPHN2 in mouse hair cells impaired balance **(a)** Schematic representation of the crossbreeding strategy to generate hair cell-specific *Lphn2* knockout mice and vestibular function analysis. **(b)** Immunostaining of LPHN2 (magenta) and Pou4f3 (green) in utricle whole mounts derived from *Pou4f3-CreER^+/-^*; *Lphn2^fl/fl^* mice (referred to as *Pc-Lphn2^fl/fl^*) or *Pou4f3-CreER^+/-^*; *Lphn2^+/+^* mice (referred to as *Pc-Lphn2^+/+^*) (n=3 mice per group). Enlarged images show the ablation of LPHN2 in the utricular hair cells of *Pc-Lphn2^fl/fl^* mice. Scale bar: 50 μm and 20 μm for low and high magnification views, respectively. **(c-f)** Quantification of the swimming scores (**c**), time on the rotating rod (**d**), travelling activity (**e**) and circling activity (**f**) in the open field test of *Pc-Lphn2^fl/fl^* mice and *Pc-Lphn2^+/+^* mice (n=11 mice per group). **(g)** Representative recording curves (left) and quantification of the VOR gain values (right) of *Pc-Lphn2^fl/fl^* mice and *Pc-Lphn2^+/+^* mice in response to earth-vertical axis rotation (n=11 mice per group). **(h)** Representative recording curves (left) and quantification of the VOR gain values (right) of *Pc-Lphn2^fl/fl^* mice and *Pc-Lphn2^+/+^* mice in response to off-vertical axis rotation (n=11 mice per group). **(i-k)** Representative click-evoked VEMP waveforms (**i**) and quantification of the P1-N1 peak amplitudes (**j**) and the P1 (filled triangle) and N1 (hollow triangle) peak latencies (**k**) of *Pc-Lphn2^fl/fl^* mice and *Pc-Lphn2^+/+^*mice at 100 dB nHL (n = 11 mice per group). (**c-h, j, k**) *P < 0.05; **P < 0.01; ***P < 0.001; ns, no significant difference. *Pc-Lphn2^fl/fl^* mice compared with *Pc-Lphn2^+/+^* mice. The bars indicate mean ± SEM values. All data were statistically analysed using unpaired two-sided Student’s *t* test.

Various behavioral tests were conducted to assess the effects of LPHN2 ablation in VHCs on balance maintenance. Notably, in addition to their significantly decreased swimming performance, the *Pou4f3-CreER^+/-^*; *Lphn2^fl/fl^*mice spent approximately 50% less time on the rotarod than their *Pou4f3-CreER^+/-^*; *Lphn2^+/+^* littermates, which was not observed in the *Lphn2^+/-^*mice (Fig. 3c, d). Moreover, compared with their *Pou4f3-CreER^+/-^*; *Lphn2^+/+^* littermates, the *Pou4f3-CreER^+/-^*; *Lphn2^fl/fl^* mice exhibited an approximately 2-fold increase in circling times in both the 2 min and 10 min testing time frames, accompanied by significantly increased traveling distances (Fig. 3e, f; Supplementary information, Fig. S5j). Furthermore, compared with the littermate control mice, the *Pou4f3-CreER^+/-^*; *Lphn2^fl/fl^*mice showed an approximately 25%∼50% decrease in the VOR response in both the earth-vertical and off-vertical axis rotation tests (Fig. 3g, h). To further evaluate the effects of LPHN2 deficiency on vestibular function, we assessed vestibular-evoked myogenic potentials (VEMPs) in the *Pou4f3-CreER^+/-^*; *Lphn2^fl/fl^* and *Pou4f3-CreER^+/-^*; *Lphn2^+/+^* mice. Whereas the positive peak (P1) and negative peak (N1) latency in the *Pou4f3-CreER^+/-^*; *Lphn2^fl/fl^*mice were not significantly altered, the P1−N1 amplitude was reduced by approximately 50% compared with that of their control littermates (Fig. 3i-k). Collectively, these results suggest that LPHN2 in VHCs plays an important role in regulating the equilibration.

To determine whether the impaired balance phenotypes in the LPHN2-deficient mice were due to defects in vestibular organ development, we further examined the morphology of utricles (and utricular hair cells) derived from the *Pou4f3-CreER^+/-^*; *Lphn2^fl/fl^* mice at P40. Our results indicated that the overall size of the utricular macula and the number of hair cells in different areas (striolar region, lateral extrastriolar region and medial extrastriolar region) of the utricle of the *Pou4f3-CreER^+/-^*; *Lphn2^fl/fl^* mice were comparable to those of their *Pou4f3-CreER^+/-^*; *Lphn2^+/+^* littermates (Fig. 4a, b). At the subcellular level, the apical surface (cuticular plate) size of the utricular hair cells, the kinocilium length and the stereocilia structure of the *Pou4f3-CreER^+/-^*; *Lphn2^fl/fl^* utricle were not significantly affected by Lphn2 ablation compared with those of their *Pou4f3-CreER^+/-^*; *Lphn2^+/+^* littermates (Fig. 4c, d; Supplementary information, Fig. S5b, c). Moreover, the expression levels and localization of MET channel components, including TMC1, TMC2, TMIE, LHFPL5 and PCDH15, in the utricular hair cells of the *Pou4f3-CreER^+/-^*; *Lphn2^fl/fl^* mice were comparable to those in the *Pou4f3-CreER^+/-^*; *Lphn2^fl/fl^*mice (Fig. 4e). These data collectively supported the normal morphology of the *Pou4f3-CreER^+/-^*; *Lphn2^fl/fl^* utricles and suggested that LPHN2 plays a regulatory role in balance sensation.

**Fig. 4.**
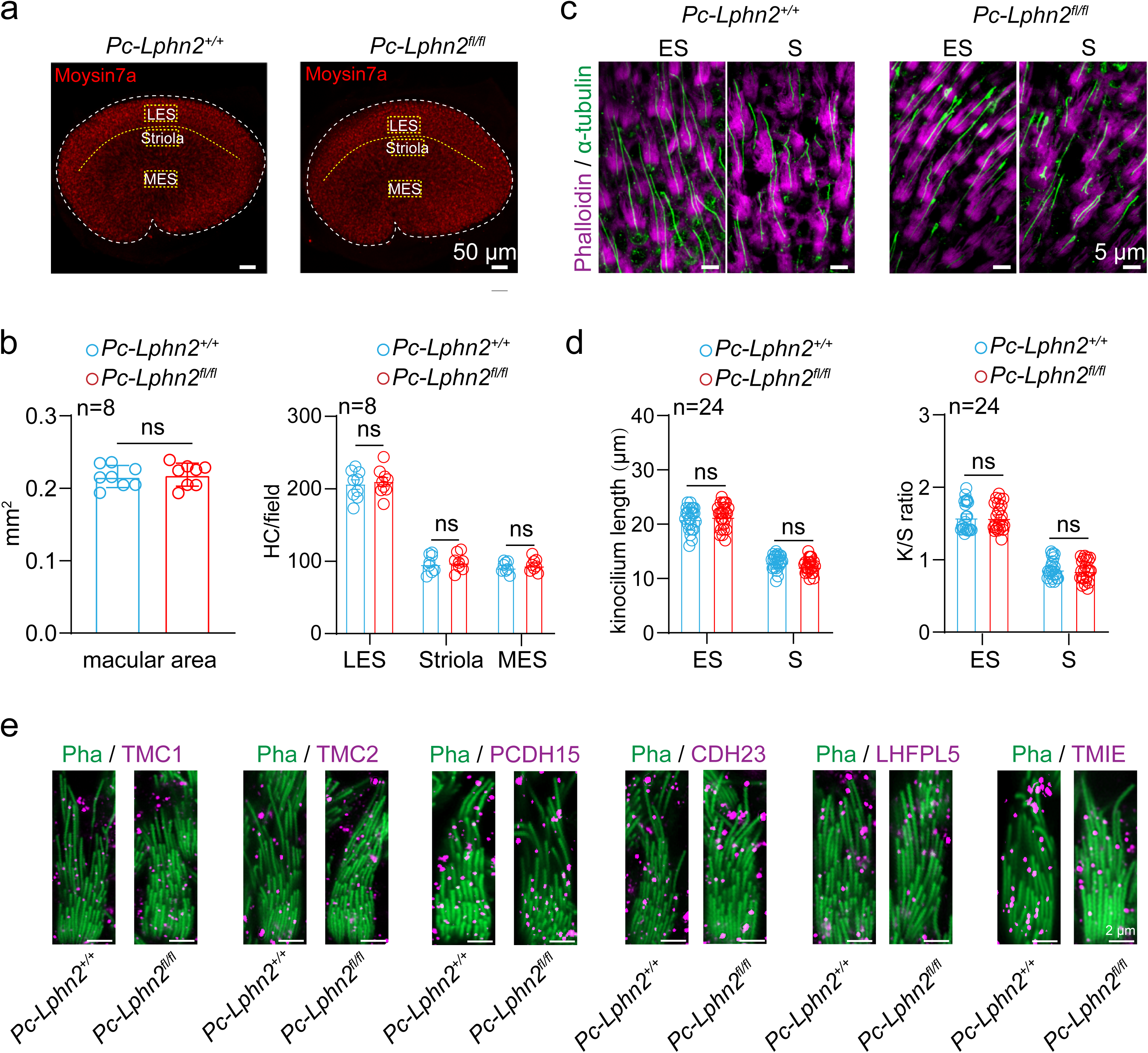
LPHN2 deficiency does not affect the morphological features of mouse utricle **(a)** Immunostaining of Myosin7a (red) in utricular hair cells derived from *Pc-Lphn2^+/+^* and *Pc-Lphn2^fl/fl^* mice at P40 (n = 8 mice per group). Scale bar, 50 μm. Three fields of 100 × 50 μm were defined and outlined in the lateral extrastriolar (LES) region, striolar region (S) and medial extrastriolar (MES) region. **(b)** Quantification of the size of utricles (left) and hair cell density at different regions of utricle (right) derived from *Pc-Lphn2^+/+^* and *Pc-Lphn2^fl/fl^* mice (n=8 mice per group). Data are correlated to Fig. 4a. **(c)** Immunostaining of kinocilium (labeled with α-tubulin, green) and stereocilia (labeled with phalloidin, magenta) in utricle whole mounts derived from *Pc-Lphn2^+/+^* and *Pc-Lphn2^fl/fl^* mice (n=3 mice per group). Scale bar, 5 μm. **(d)** Quantification of the length of kinocilium (left) and the ratio of lengths of the kinocilium to tallest stereocilia (right) in ES or S region of utricle whole mounts derived from *Pc-Lphn2^+/+^*and *Pc-Lphn2^fl/fl^* mice. Data are correlated to Fig. 4c (n = 24 hair cells from 3 mice per group). **(e)** Co-immunostaining of phalloidin (referred to as Pha, green) and different MET machinery components (magenta), including TMC1, TMC2, PCDH15, CDH23, LHFPL5 and TMIE, in utricular hair cells derived from *Pc-Lphn2^+/+^* and *Pc-Lphn2^fl/fl^* mice (n = 6 mice per group). **(b, d)** ns, no significant difference. *Pc-Lphn2^+/+^* mice compared with *Pc-Lphn2^fl/fl^* mice. The bars indicate mean ± SEM values. All data were statistically analyzed using unpaired two-sided Student’s *t* test.

### Genetic deficiency or pharmacological blockade of LPHN2 impairs the utricle MET current

To investigate whether LPHN2 participates in equilibrioception, we examined MET responses using isolated utricles. We employed a fluid jet system to deflect hair bundles in utricular hair cells and recorded the corresponding MET currents using a whole-cell patch-clamp technique^7,52,53^. To specifically record the MET response in the LPHN2-expressing utricular hair cells, we labeled these cells by using a modified AAV-ie-*Lphn2pr*-mCherry vector, which enabled the expression of the fluorescent protein mCherry driven by the *Lphn2* promoter (Fig. 5a; Supplementary information, Fig. S6a, b). AAV-ie-*Lphn2pr*-mCherry was injected into P3 mice through a round window membrane, and the mCherry-labeled utricular hair cells at P10 were selected for fluid jet stimulation and MET recording at a holding potential of -64 mV^54^. Notably, the peak MET currents in the mCherry-labeled utricular hair cells (from the medial extrastriolar region) of the *Pou4f3-CreER^+/-^*; *Lphn2^fl/fl^* mice were reduced by approximately 50% compared with those from the control *Pou4f3-CreER^+/-^*; *Lphn2^+/+^* littermates (225.8 ± 12.4 pA v.s. 112.4 ±7.4 pA), suggesting that LPHN2 is involved in the MET process in utricular hair cells (Fig. 5b, c).

**Fig. 5.**
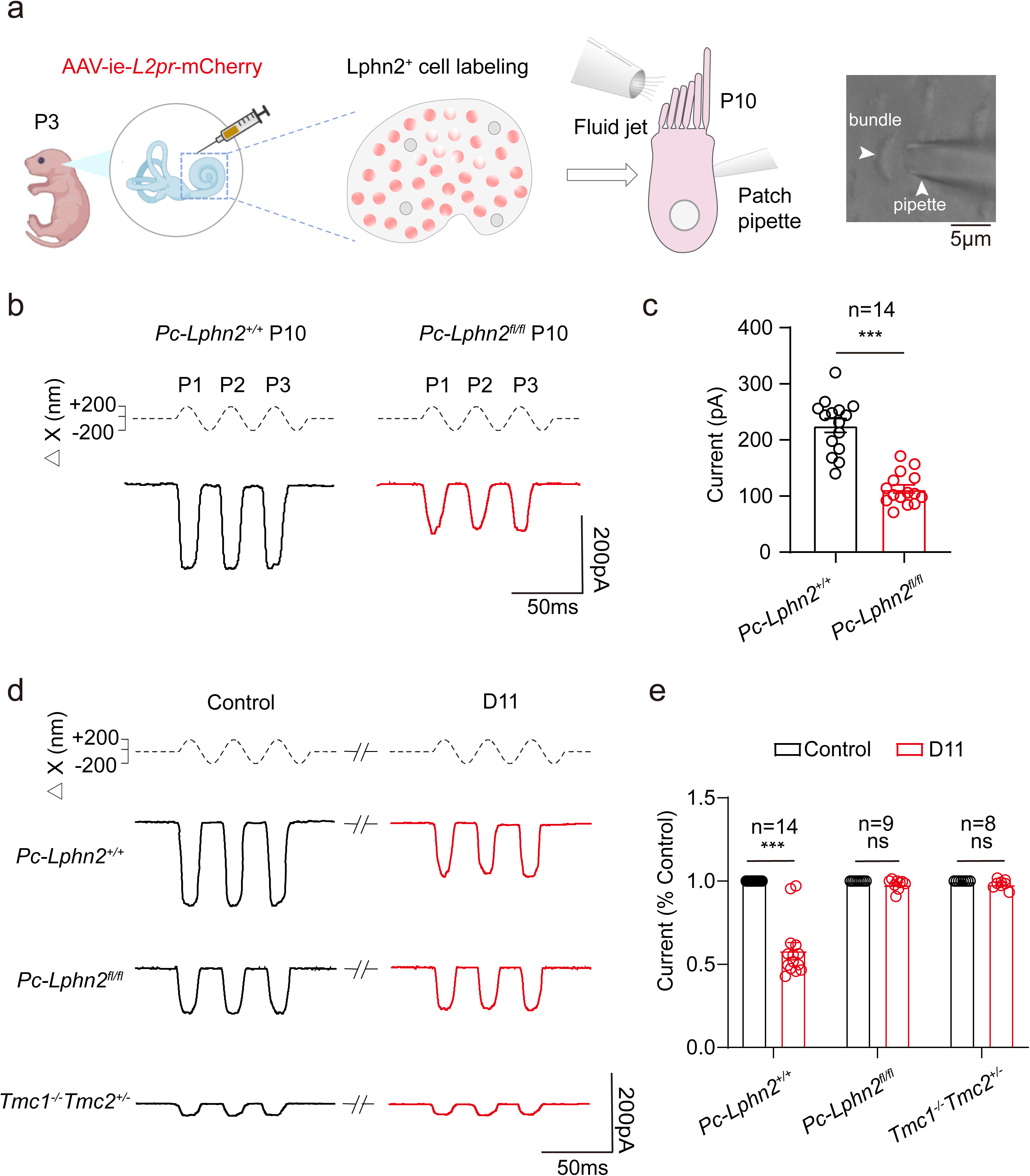
LPHN2 deficiency impairs MET currents in utricular hair cells **(a)** Schematic illustration of the labelling of LPHN2-expressing utricular hair cells by AAV-ie-*Lphn2pr*-mCherry (referred to as AAV-ie-*L2pr*-mCherry) and the MET current recording by fluid-jet stimulation. **(b)** Representative MET current traces induced by sinusoidal fluid-jet stimulation in utricular hair cells of *Pc-Lphn2^+/+^* mice (black) or *Pc-Lphn2^fl/fl^*mice (red) at P10. **(c)** Quantification of the saturated MET currents in utricular hair cells of *Pc-Lphn2^+/+^* mice or *Pc-Lphn2^fl/fl^* mice at P10 (n = 14). Data are correlated to Fig. 5b. **(d, e)** Representative current traces (c) and quantitative analysis (e) of fluid jet-stimulated MET responses in utricular hair cells derived from *Pc-Lphn2^+/+^*, *Pc-Lphn2^fl/fl^* or *Tmc1^-/-^Tmc2*^+/-^mice in the absence (black) or presence (red) of 50 nM D11. Data are normalized to the saturated MET response of control vehicle-treated hair cells in respective groups (n = 14, 9 and 8 for *Pc-Lphn2^+/+^*, *Pc-Lphn2^fl/fl^*and *Tmc1^-/-^* respectively). Data are correlated to Supplementary information, Fig.S6f. **(c)** ***P < 0.001; *Pc-Lphn2^+/+^* mice compared with *Pc-Lphn2^fl/fl^*mice. **(e)** ***P < 0.001; ns, no significant difference. Utricular hair cells treated with D11 compared with those treated with control vehicle. The bars indicate mean ± SEM values. All data were statistically analysed using unpaired (**c**) or paired (**e**) two-sided Student’s *t* test.

To further investigate the regulatory role of LPHN2 in MET and to exclude the possibility that dampened MET in LPHN2-deficient mice might be caused by potential deficits in certain MET machinery components, we next utilized a reversible and selective inhibitor of LPHN2, named D11 (developed by structure-based in silico screening and described in the companion manuscript), to investigate the regulatory mechanism of LPHN2 in utricular MET (Supplementary information, Fig. S6c). Consistent with the MET data obtained in the Lphn2-deficient utricular hair cells, pretreatment of utricular explants derived from the *Pou4f3-CreER^+/-^*; *Lphn2^+/+^* mice with 50 nM D11 caused an approximately 43% decrease in the MET current, which returned to the normal level after D11 was removed (Fig. 5d, e; Supplementary information, Fig. S6d-f). As a negative control, the inhibitory effects of D11 on MET currents in LPHN2 promoter-labeled VHCs were abolished in utricular hair cells derived from the *Pou4f3-CreER^+/-^*; *Lphn2^fl/fl^* mice, suggesting a specific role for LPHN2 in MET regulation. Intriguingly, the residual MET response in utricular hair cells derived from *Tmc1^-/-^Tmc2*^+/-^ mice, which might be regulated by TMC2 compensation or by other unknown channels, was not significantly altered by D11 administration (Fig. 5d, e; Supplementary information, Fig. S6f). These data collectively suggested that mechanosensitive LPHN2 actively participates in the MET process in utricular hair cells, potentially through crosstalk with TMC1, the ion-conducting pore of the MET channel complex.

### LPHN2 regulates the MET current at the apical surface of utricular hair cells

Notably, we observed that LPHN2 is expressed only at the apical surface and not at the stereocilia of utricular hair cells, suggesting that LPHN2 modulates MET in a tip-link-independent manner. Previous studies have indicated that whereas the outward phase of the sinusoidal flow in the fluid jet assay induces a normal-polarity MET current by deflecting the hair bundle toward the longest stereocilia in mature cochlear or VHCs, the inward fluid flow in the opposite direction can evoke a reverse-polarity current in hair cells lacking tip links or key MET channel components (e.g., *Tmc1^-/-^Tmc2*^-/-^ or *Tmie*^-/-^)^52,53,55^. Following previously established protocol, we treated the hair cells of utricles with the calcium-chelating agent BAPTA to disrupt tip links and investigated the potential role of LPHN2 in tip-link-independent MET in utricles (Fig. 6a). Consistent with previous reports, after cochlear explants were treated with BAPTA for 5 min, the normal-polarity MET currents (782.5 ± 15.0 pA) were replaced by reverse-polarity currents (439.3 ± 26.0 pA) in response to sinusoidal fluid jet stimulation (Fig. 6b, c)^52,53^. Unexpectedly, treatment of utricular explants with BAPTA under identical conditions to those of the cochlear explants resulted in unique biphasic MET currents, including a reduced normal-polarity MET current (48.2% decrease compared with the control normal-polarity current) and a newly emerging reverse-polarity current (Fig. 6b, c). These data suggested the coexistence of tip-link-independent normal-polarity and reverse-polarity MET currents in utricular hair cells, which was different from the MET characteristics of cochlear hair cells (CHCs).

**Fig. 6.**
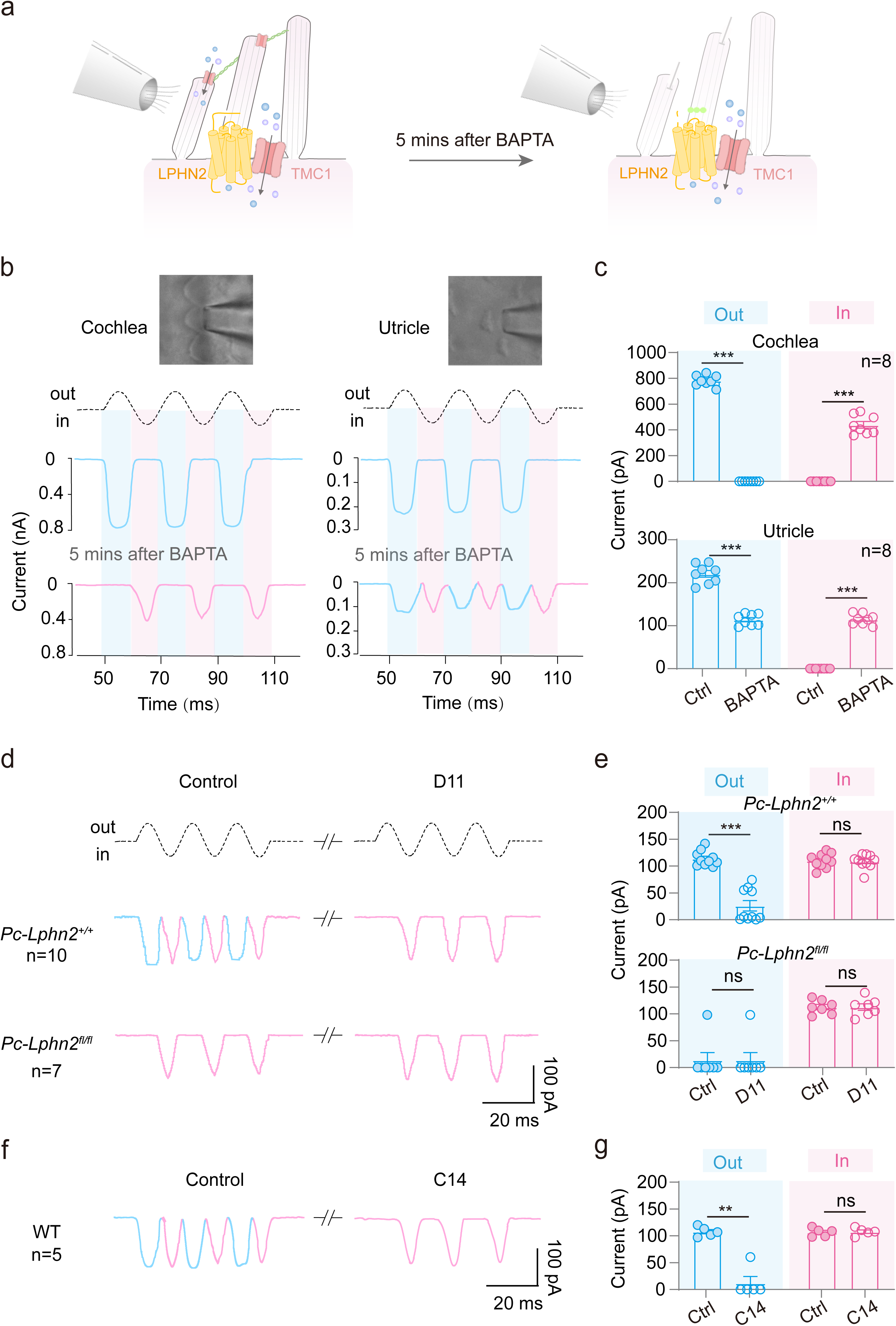
LPHN2 regulates the MET current at the apical surface of utricular hair cells (a) Schematic illustration showing the fluid jet-stimulated MET responses in utricular hair cells before and after treatment with BAPTA (b) Representative MET current traces induced by sinusoidal fluid-jet stimulation in cochlear OHCs (left) or utricular hair cells (right) before and after treatment with BAPTA for 5 min (n = 8 per group). The normal-polarity and reverse-polarity MET current traces are colored blue and pink, respectively. (c) Quantification of the saturated normal-polarity (outward phase, blue) or reverse-polarity (inward phase, pink) MET current of cochlear (top) or utricular hair cells (bottom) in response to fluid-jet stimulation (n = 8 per group). **(d, e)** Representative current traces (**d**) and quantitative analysis (**e**) of fluid jet-stimulated MET responses in BAPTA-treated utricular hair cells derived from *Pc-Lphn2^+/+^* or *Pc-Lphn2^fl/fl^* mice in the absence or presence of 50 nM D11 (n = 10 and 7 for *Pc-Lphn2^+/+^* and *Pc-Lphn2^fl/fl^* mice, respectively). **(f, g)** Representative current traces (**f**) and quantitative analysis (**g**) of fluid jet-stimulated MET responses in BAPTA-treated WT utricular hair cells in the absence or presence of 1 μM C14 (n=5). **(c)** ***P < 0.001; Hair cells pretreated with BAPTA compared with those treated with control vehicle. **(e, g)** **P < 0.01; ***P < 0.001; ns, no significant difference. Utricular hair cells treated with D11 or C14 compared with those treated with control vehicle. The bars indicate mean ± SEM values. All data were statistically analysed using paired two-sided Student’s *t* test.

Specifically, the residual normal-polarity MET current recorded in the BAPTA-treated utricular hair cells derived from the WT or *Pou4f3-CreER^+/-^*; *Lphn2^+/+^* mice was absent in those derived from the *Pou4f3-CreER^+/-^*; *Lphn2^fl/fl^* mice (Fig. 6d, e). Therefore, we speculated that LPHN2 expressed at the apical surface of utricular hair cells might be responsible for tip-link-independent normal-polarity MET currents. Consistent with this hypothesis, pretreatment with the LPHN2-specific inhibitor D11 (50 nM) abrogated the residual normal-polarity MET current in the BAPTA-treated utricular hair cells derived from the *Pou4f3-CreER^+/-^*; *Lphn2^+/+^* mice, which recovered to normal levels after D11 was washed out (Fig. 6d, e; Supplementary information, Fig. S6g, h). In contrast, the reverse-polarity currents were not significantly affected by D11 treatment in either *Pou4f3-CreER^+/-^*; *Lphn2^+/+^*or *Pou4f3-CreER^+/-^*; *Lphn2^fl/fl^* utricular hair cells (Fig. 6d, e). To further determine the potential roles of TMC1 in tip link-independent MET currents, we developed a reversible inhibitor through structure-based in silico screening (a simulated structure of mouse TMC1 was modeled using Alphafold2) (Supplementary information, Fig. S7a-d); this inhibitor C14 showed inhibitory effects on TMC1, but not on several other ion channels, such as CNG channel *in vitro* or sodium channels in primary utricular hair cells (Supplementary information, Fig. S7e-g). We revealed that, similar to LPHN2, pharmacological inhibition of TMC1 by the inhibitor also abolished the residual normal-polarity currents without affecting the reverse-polarity currents (Fig. 6f, g; Supplementary information, Fig. S7h). Collectively, these data indicated that LPHN2 plays an important role in regulating previously uncharacterized normal-polarity MET currents at the apical surface of utricular hair cells, potentially by interacting with TMC1.

### Colocalization and functional coupling of LPHN2 with TMC1 at the apical surface

In our companion manuscript, by performing proteomic interactome analyses using purified LPHN2 as bait, as well as the co-immunostaining assay, our data suggested a direct interaction between LPHN2 and TMC1 in the mouse cochlea ^56^. A similar interaction between LPHN2 and TMC1 was also detected by in vivo coimmunoprecipitation of mouse utricular lysates (Fig. 7a). Several studies have shown that TMC1 is distributed at the apical surface of hair cells^9,57,58^. We observed fluorescent puncta of TMC1 at the apical surface of the utricular hair cells by optical sectioning microscopy, where TMC1 was significantly coimmunostained with LPHN2, suggesting the potential localized assembly of these two membrane proteins in vivo (Fig. 7b).

**Fig. 7.**
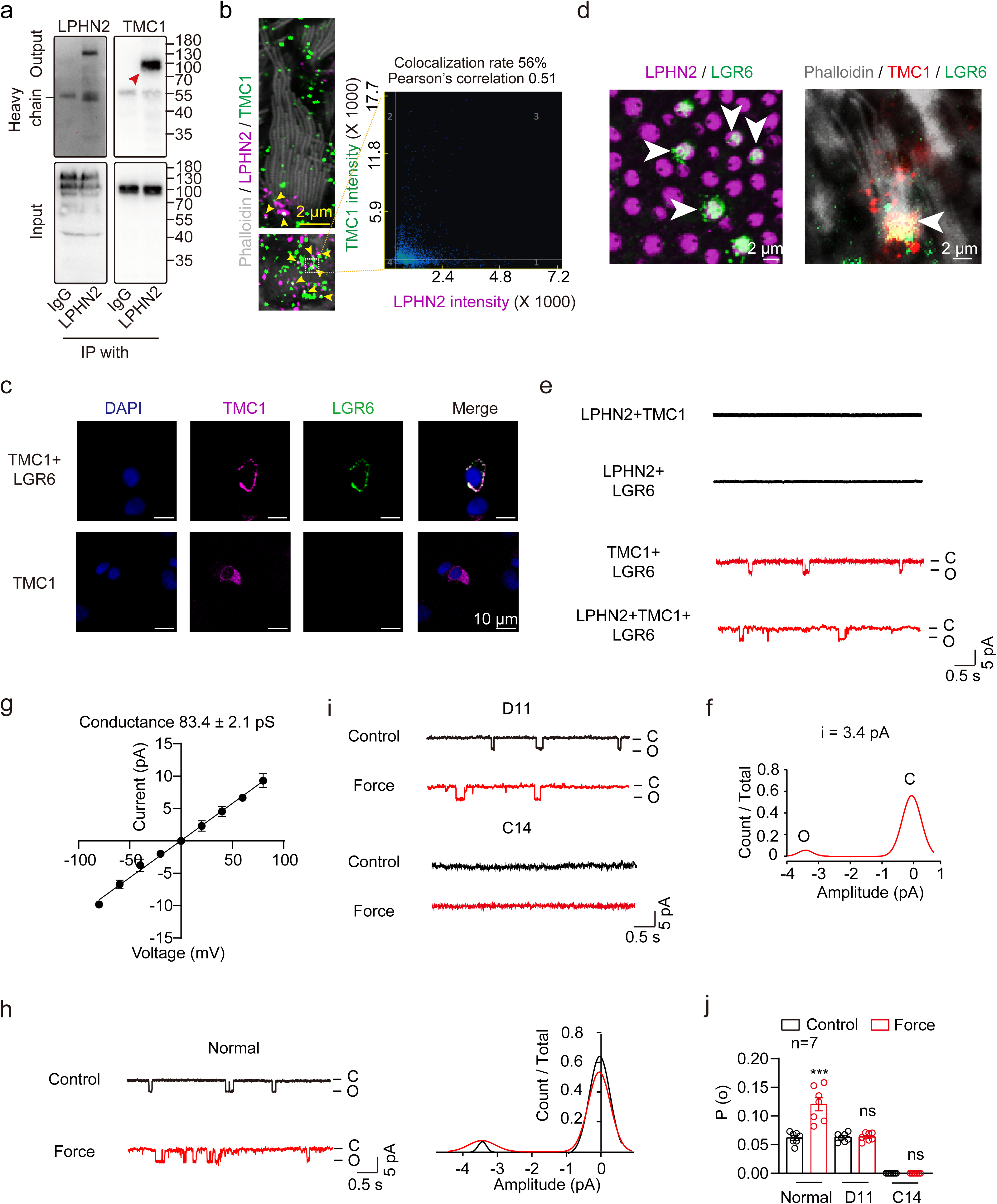
LPHN2 colocalizes with TMC1 at the apical surface of utricular hair cells and regulates MET through coupling to TMC1 **(a)** Co-immunoprecipitation of LPHN2 with TMC1 in the lysates of mouse utricles. Representative blotting from three independent experiments is shown (n = 3). **(b)** Co-immunostaining of LPHN2 (magenta) with TMC1 (green) in the apical surface of utricular hair cells in utricle whole mounts derived from WT mice (n = 6 mice per group). Scale bar: 2 μm. Arrows indicate the colocalization of LPHN2 with TMC1 at the apical surface of utricular hair cells. **(c)** Co-immunostaining of TMC1 (magenta) with LGR6 (green) in HEK293 cells transfected with TMC1 only or with TMC1 and LGR6. Scale bar: 10 μm. Representative images from three independent experiments were shown (n = 3). **(d)** Co-immunostaining of LGR6 (green) with LPHN2 (magenta) or with TMC1 (red) at the apical surface of utricular hair cells. Scale bar, 2 μm. Representative images from three independent experiments were shown (n = 3). **(e)** Representative spontaneous single-channel currents of TMC1 at -40 mV recorded in HEK293 cells co-transfected with LPHN2/TMC1, LPHN2/LGR6, TMC1/LGR6 or LPHN2/TMC1/LGR6. Representative current traces from three independent experiments were shown (n=3). **(f)** The normalized all-point amplitude histogram analysis of single-channel currents in HEK293 cells transfected with TMC1/LGR6. The distribution data were fitted by a sum of two Gaussians, and the peaks correspond to the closed (C) and open (O) states. **(g)** The current-voltage (I-V) relationship of the spontaneous currents recorded in HEK293 cells transfected with TMC1/LGR6 (n = 3). **(h)** Representative current traces (left) and histogram analysis (right) of the single-channel currents recorded in HEK293 cells transfected with LPHN2/TMC1/LGR6 under control condition (black) or in response to 10 pN force stimulation (red). **(i)** Representative spontaneous and force (10 pN)-stimulated single-channel current traces recorded in HEK293 cells transfected with LPHN2/TMC1/LGR6 in the presence of 50 nM D11 (top) or 1 μM C14 (bottom). **(j)** Effect of D11 or C14 treatment on the force-induced single-channel open probability (n = 7). Data are correlated to Fig. 7h, i. (**j**) ***p < 0.001; ns, no significant difference. Force-stimulated cells compared with control cells. The bars indicate mean ± SEM values. All data were statistically analysed using paired two-sided Student’s *t* test.

To further delineate the interaction between LPHN2 and TMC1, we next attempted to reconstitute the functional coupling between LPHN2 and TMC1 in a heterologous system. We previously identified two GPCRs that are endogenously expressed in hair cells and can transport TMC1 to the plasma membrane in HEK293 cells (described in our companion paper ^56^) (Fig. 7c). One of these two receptors, LGR6, was found to be expressed at the apical surface of utricle hair cells, where it coimmunostained with LPHN2 and TMC1 (Fig. 7d). Therefore, we selected this receptor to function as the trafficking chaperone for TMC1 in the heterologous system, which we reasoned would more closely mimic the endogenous landscape. By performing patch-clamp recording at a holding potential of -40 mV, we observed comparable spontaneous single-channel opening in the HEK293 cells cotransfected with TMC1/LPHN2/ LGR6 and the cells cotransfected with TMC1/LGR6 but not in the negative control cells cotransfected with TMC1/LPHN2 or with LPHN2/ LGR6 (Fig. 7e). The average single-channel TMC1 current and conductance were 3.4 pA and 83.4 ± 2.1 pS, respectively, suggesting the successful trafficking of TMC1 onto the plasma membrane by LGR6 (Fig. 7f, g). Moreover, the application of 10 pN force through LPHN2-M-beads to the heterologous system coexpressing LPHN2/TMC1/LGR6 induced an approximately 1.9-fold increase in the probability of TMC1 opening compared with the resting condition, which was abrogated by pretreatment with the LPHN2-specific inhibitor D11 or with the TMC1 inhibitor C14 (Fig. 7h-j). Therefore, these results indicated LPHN2 can sense extracellular mechanical stimuli to enhance the TMC1 activity.

### Force sensation by LPHN2 induces neurotransmitter release and Ca^2+^ influx in utricular hair cells

In response to mechanical stimulation in equilibrioception, the VHCs may release neurotransmitters, such as glutamate, to transmit positional or motional information to spiral ganglion neurons (SGNs). We therefore specifically expressed the genetically encoded glutamate sensor R^ncp^-iGluSnFR (R^ncp^-iGlu) in SGNs to monitor real-time glutamate release from utricular hair cells^59^. Binding of glutamate to the R^ncp^-iGlu sensor induced conformational-based modulation of the protonation state of the chromophore, leading to a reduction in fluorescence intensity (Fig. 8a). Notably, repeated force application with 10 pN on the utricular sensory epithelium through LPHN2-M-beads induced detectable glutamate release, which disappeared with force removal (Fig. 8b, c). The magnitude of the LPHN2-M-beads-induced glutamate release from utricular hair cells was approximately half of that induced by force applied on tip link component CDH23 via CDH23-M-beads, which served as a positive control (Supplementary information, Fig. S8a, b). As a negative control, the force applied by the Ctrl-beads did not induce any detectable glutamate secretion (Fig. 8b, c; Supplementary information, Fig. S8a, b). Moreover, force-stimulated glutamate secretion via LPHN2-M-beads was abrogated in the WT utricular explants pretreated with the LPHN2-specific inhibitor D11 or TMC1 inhibitor C14 or in the utricular explants derived from the *Pou4f3-CreER^+/-^*; *Lphn2^fl/fl^* mice (Fig. 8d; Supplementary information, Fig. S8c, d). These results supported a modulatory role of LPHN2 in converting mechanical stimuli into glutamate release.

**Fig. 8.**
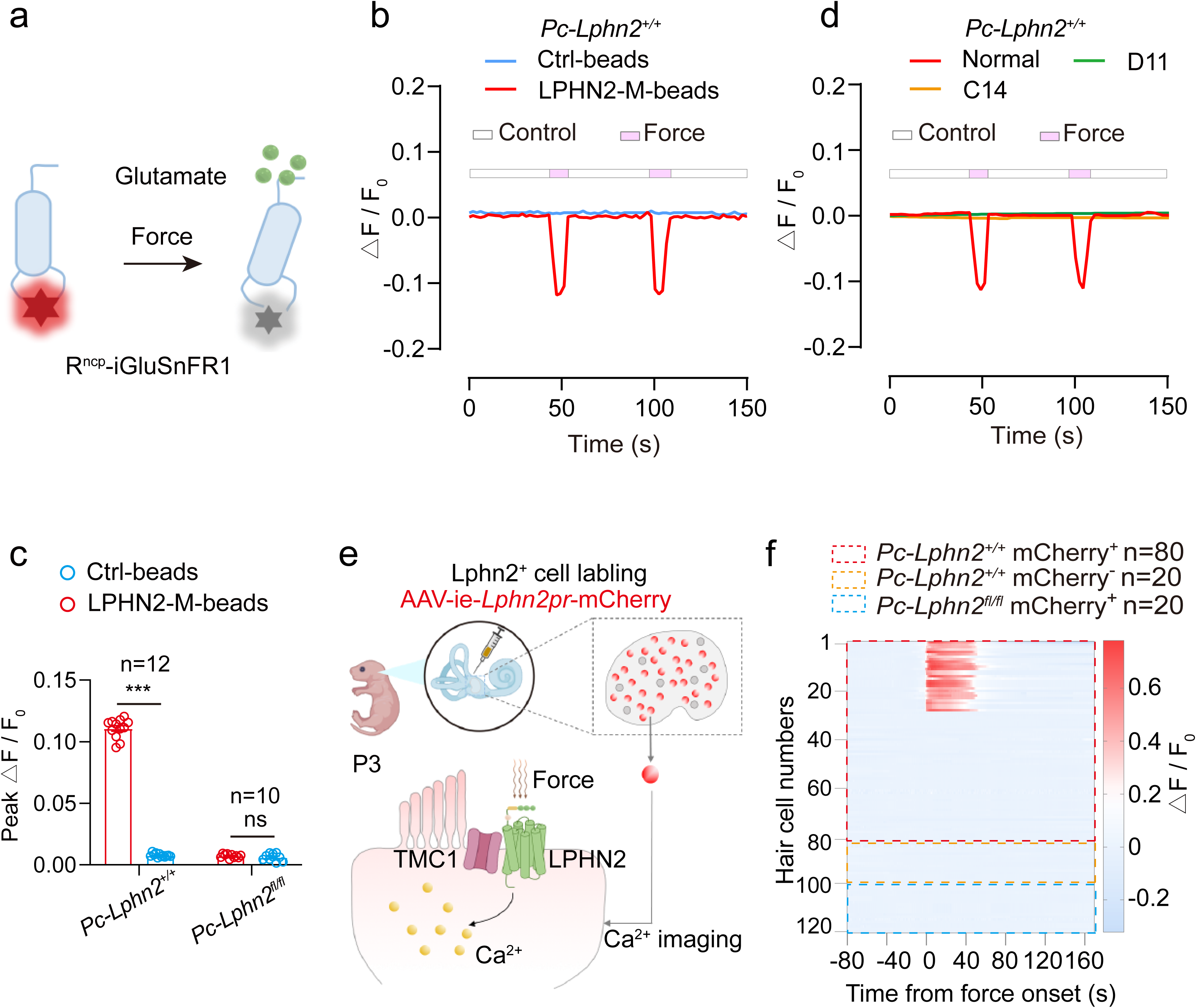
Force sensation by LPHN2 induces glutamate release and Ca^2+^ signals in utricular hair cells **(a)** Schematic diagram showing the working mechanism of the red fluorescent glutamate sensor Rncp-iGluSnFR (Rncp-iGlu). **(b, c)** Representative traces (**b**) and quantitative analysis (**c**) of glutamate secretion from individual utricular hair cell derived from P10 *Pc-Lphn2^+/+^* mice (n=12) or *Pc-Lphn2^fl/fl^* mice (n=10) in response to force applied by LPHN2-M-beads or control beads. Data are correlated to Supplementary information, Fig. S8c. **(d)** Representative traces of glutamate secretion from individual utricular hair cell derived from P10 *Pc-Lphn2^+/+^*mice pretreated with control vehicle, 50 nM D11 or 1 μM C14 in response to force applied by LPHN2-M-beads (n = 12). Data are correlated to Supplementary information, Fig. S8d. **(e)** Schematic illustration of Ca^2+^ imaging in mouse utricular hair cells labelled with AAV-ie-*L2pr*-mCherry in response to force stimulation. **(f)** Heatmaps showing the Ca^2+^ responses in individual utricular hair cell derived from *Pc-Lphn2^+/+^* or *Pc-Lphn2^fl/fl^* mice. n=80, 20 and 20 for mCherry-labelled *Pc-Lphn2^+/+^* cells (red box), mCherry-unlabelled *Pc-Lphn2^+/+^* cells (orange box), and mCherry-labelled *Pc-Lphn2^fl/fl^* cells (blue box), respectively. The color intensity represents the magnitude of the calcium response characterized by △F/F_0_. **(c)** ***P < 0.001; ns, no significant difference. Utricle explants treated with LPHN2-M-beads compared with those treated with Ctrl-beads. The bars indicate mean ± SEM values. All data were statistically analysed using unpaired two-sided Student’s *t* test.

Intracellular Ca^2+^ is a key element for neurotransmitter release and may play a pivotal role in equilibration sensation^60–63^. We next examined the Ca^2+^ response downstream of LPHN2 in single hair cells after force stimulation. The Ca^2+^ signal in the LPHN2-expressing utricular hair cells, which were labeled with AAV-ie-*Lphn2pr*-mCherry, in response to force stimulation was recorded by a Fura-2 fluorescent probe. Approximately 35% of the LPHN2-expressing hair cells derived from the *Pou4f3-CreER^+/-^*; *Lphn2^+/+^*utricle showed an increase in Ca^2+^ levels in response to force stimulation (Fig. 8e, f). In contrast, the AAV-ie-*Lphn2pr*-mCherry-labeled utricular hair cells derived from the *Pou4f3-CreER^+/-^*; *Lphn2^fl/fl^* mice showed no detectable Ca^2+^ signals in response to force application via LPHN2-M beads (Fig. 8f). We speculated that the relatively lower response rate of the LPHN2-expressing hair cells in the Ca^2+^ assay compared to that in the fluid jet assay (∼86%, 12 out of 14 cells were sensitive to D11 treatment) might be due to the inefficient interaction between the LPHN2-M-beads and the endogenous LPHN2 (Fig. 5d, e). Consistent with this hypothesis, only about 53% of the randomly-selected utricular hair cells showed detectable Ca^2+^ signals in response to mechanical stimulation applied on tip link component CDH23, suggesting that binding efficiency of the magnetic beads was around 50% (Supplementary information, Fig. S8e, f).

Taken together, our results collectively indicate that the activation of LPHN2 by force stimulation induces Ca^2+^ signaling and neurotransmitter release in VHCs.

### Re-expression of LPHN2 in VHCs rescued the equilibrioception of LPHN2-deficient mice

To further assess the functional roles of LPHN2 specifically expressed in the vestibular system, we reintroduced LPHN2 expression in the utricular hair cells of the *Pou4f3-CreER^+/-^*; *Lphn2^fl/fl^* mice via AAV delivery. Our previous structural and functional analyses revealed that force sensation by adhesion GPCRs was primarily mediated by the juxtamembrane GAIN domain^18,20^. The Flag-tagged LPHN2-GAIN construct with the deletion of 523 N-terminal residues retained the mechanosensitive potential and showed dose-dependent Gs signaling in response to force stimulation with Flag-M-beads, and the response was comparable to that of full-length LPHN2 when expressed at similar levels (described in the companion paper ^56^). Therefore, we packaged the Flag-LPHN2-GAIN sequences into the AAV-ie-*Lphn2pr*-mCherry vector (mCherry following an IRES element was fused to the C-terminus of LPHN2-GAIN, referred to as AAV-ie-LPHN2) and delivered the virus into P3 *Pou4f3-CreER^+/-^*; *Lphn2^fl/fl^* mice through a round window membrane injection using AAV-ie-*Lphn2pr*-mCherry as a negative control (Supplementary information, Fig. S9a). LPHN2-GAIN was specifically expressed in the utricular hair cells, but not in the brains, of the *Pou4f3-CreER^+/-^*; *Lphn2^fl/fl^* mice 14 days after the administration of AAV-ie-LPHN2, as shown by western blotting and immunofluorescence analysis (Supplementary information, Fig. S9b, c). As a negative control, the *Pou4f3-CreER^+/-^*; *Lphn2^fl/fl^* mice infected with AAV-ie-*Lphn2pr*-mCherry exhibited only specific utricular hair cell labeling but no detectable LPHN2-GAIN expression (Supplementary information, Fig. S9b, c).

We next assessed the equilibration-related behaviors of the mice receiving LPHN2-GAIN gene delivery. Notably, the LPHN2-deficient mice treated with AAV-ie-LPHN2 exhibited significantly improved performance in all behavior studies related to equilibrioception, including the forced swimming, open field and rotarod tests, and the performances were comparable to those of the WT mice (Fig. 9a-d). Moreover, the VOR gain values of the AAV-ie-LPHN2-treated *Pou4f3-CreER^+/-^*; *Lphn2^fl/fl^* mice at all tested frequencies in both the earth-vertical and off-vertical axis rotation tests recovered to levels comparable to those of the WT mice (Fig 9e, f). As a negative control, the LPHN2-deficient mice treated with AAV-ie-*Lphn2pr*-mCherry showed no significant improvement in performance in any of the above tests (Fig. 9a-f). By analyzing the MET response of the utricular hair cells, we found that *Lphn2*-deficient utricular hair cells infected with AAV-ie-LPHN2, but not those infected with AAV-ie-*Lphn2pr*-mCherry, showed significantly increased MET currents and restored D11 responsiveness, which were comparable to those of the WT utricular hair cells (Supplementary information, Fig S9d, e). Furthermore, after BAPTA treatment to disrupt the tip-links, the AAV-ie-LPHN2-treated *Pou4f3-CreER^+/-^*; *Lphn2^fl/fl^* utricular hair cells reproduced a normal-polarity MET current, which was comparable to that found in the BAPTA-treated WT utricular hair cells and could be ablated by D11 administration (Fig. 9g, h). In contrast, the *Pou4f3-CreER^+/-^*; *Lphn2^fl/fl^* vestibular hair cells treated with AAV-ie-*Lphn2pr*-mCherry showed no significant alterations in MET characteristics (Fig. 9g, h). Therefore, reintroduction of mechanosensitive LPHN2-GAIN specifically in utricular hair cells rescued equilibrioception and restore D11-sensitive MET in LPHN2-deficient mice. These findings further indicated that force-activated LPHN2 in the vestibular system directly contributes to normal equilibrioception.

**Fig. 9.**
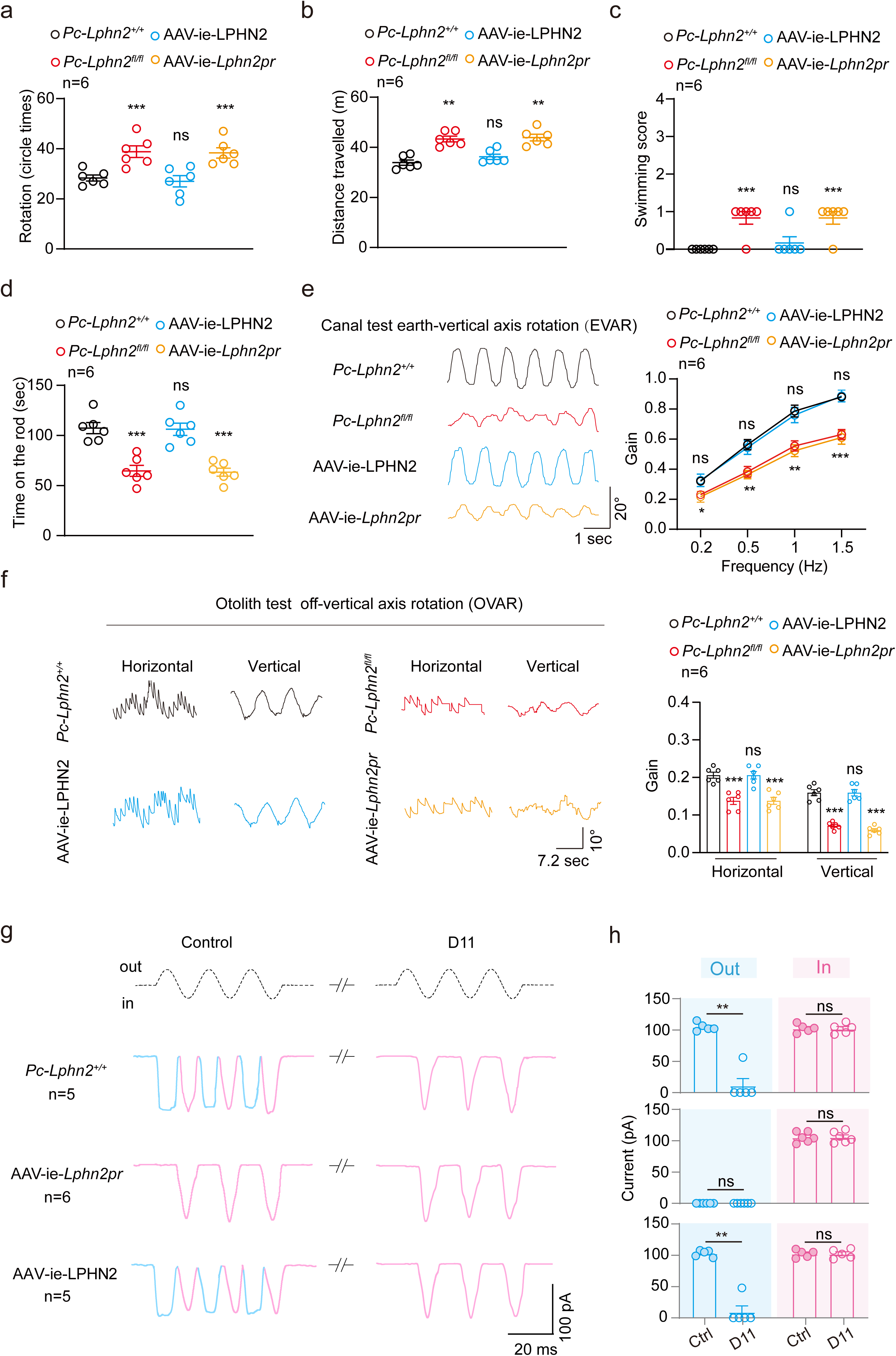
Specific re-expression of LPHN2 in hair cells rescued the vestibular function of LPHN2-deficient mice **(a-d)** Quantification of the circling (**a**) and travelling activity (**b**) in the open-field test during 10 min, swimming scores (**c**) and duration time on the rotating rod (**d**) of *Pc-Lphn2^+/+^*, *Pc-Lphn2^fl/fl^*, AAV-ie-LPHN2 mice and AAV-ie-*Lphn2pr* mice (n=6 mice per group). **(e, f)** Representative recording curves (left) and quantification of the VOR gain responses (right) of *Pc-Lphn2^+/+^*, *Pc-Lphn2^fl/fl^*, AAV-ie-LPHN2 mice and AAV-ie-*Lphn2pr* mice to earth-vertical axis (**e**) or off-vertical axis (**f**) rotation (n=6 mice per group). **(g,h)** Representative current traces (**g**) and quantitative analysis (**h**) of fluid jet-stimulated MET responses in BAPTA-treated utricular hair cells derived from *Pc-Lphn2^+/+^*, AAV-ie-*L2pr-*LPHN2 or AAV-ie-*L2pr-*mCherry mice in the absence or presence of 50 nM D11 (n = 5, 6 and 5 for *Pc-Lphn2^+/+^*, AAV-ie-*L2pr-*LPHN2 or AAV-ie-*L2pr-*mCherry mice, respectively). (**a-f**) *P < 0.05; **P < 0.01; ***P < 0.001; ns, no significant difference. *Pc-Lphn2^fl/fl^*, AAV-ie-*L2pr-*LPHN2 or AAV-ie-*L2pr-*mCherry mice compared with *Pc-Lphn2^+/+^* mice. (**h**) **P < 0.01; ns, no significant difference. Utricular hair cells treated with D11 compared with those treated with control vehicle. The bars indicate mean ± SEM values. All data were statistically analysed using one-way ANOVA with Dunnett’s post hoc test (**a-f**) or paired two-sided Student’s *t* test (**h**).

## Discussion

Despite its importance for daily life and motion in three-dimensional space, the molecular mechanism underlying the sense of balance is not fully understood. Studies have revealed that ion channels, such as TMC1/2, are potential key components of the MET process, which is essential for equilibrioception^12,64,65^. In addition to ion channels, another group of membrane receptors, GPCRs, are known as direct sensors for vision, odor and touch^29,37,66^. Although recent studies have shown that some GPCR members, such as CELSR1, GPP156 and GPR126, contribute to vestibular development or the maintenance of VHC planar polarity^67–69^, GPCRs are generally excluded from the equilibrioception process in the peripheral vestibular system due to their relatively slow kinetics in mediating secondary messenger pathways. In our current study, by screening potential mechanosensitive aGPCRs in the vestibular system, we revealed that the force-sensing protein LPHN2 plays an important role in maintenance of normal balance. Importantly, whereas conditional KO of LPHN2 expression in VHCs impaired the balance behaviors of mice without affecting the morphology of the utricular macula or hair cells, specific re-expression of LPHN2 in VHCs of the Lphn2-deficient mice rescued equilibrioception. Specifically, both tamoxifen treatment and AAV delivery in the *Pou4f3-CreER^+/-^*; *Lphn2^fl/fl^* mice were limited to the vestibular organs through round window membrane injection, which resulted in no detectable effects on LPHN2 expression in tissues beyond the inner ear, especially the vestibular or somatosensory nuclei in the brainstem. These results collectively suggested a specific role for LPHN2 in regulating balance sensation in the peripheral vestibular system.

The LPHN2 actively participated in the MET process in VHCs, which was supported by the results of a fluid jet assay using VHCs derived from Lphn2-deficient mice or a specific reversible LPHN2 inhibitor. Unexpectedly, in contrast to the unique presence of LPHN2 near the tips of stereocilia in CHCs^56^, LPHN2 was absent in stereocilia and was only expressed at the apical surface of utricular hair cells. Importantly, through local interactions with TMC1, LPHN2 at the apical surface regulates a tip link-independent normal-polarity MET current, which contributes to approximately 50% of the total MET current in response to fluid jet stimulation. Therefore, our findings revealed a previously uncharacterized MET process at the apical surface of VHCs mediated by a GPCR-TMC1 functional coupling pair. Notably, previous studies have reported nonidentical MET properties, including current amplitude and adaptation parameters, in utricular hair cells stimulated with a fluid jet (mechanical stimulation of both the hair bundle and apical surface) and those deflected with a stiff probe (stimulation of the hair bundle only), with potentially unknown underlying mechanisms^54,70^.

Our findings herein have thus provided a possible explanation for this discrepancy. Moreover, a tip link-independent reverse-polarity MET current has been previously identified in CHCs; this current was reported to be regulated by Piezo2 and could be evoked following disruption of the sensory-transduction machinery^52,71^. A similar reverse-polarity MET current was also present in the saccular hair cells of mice during the embryonic period but disappeared when normal MET currents developed^53^. Our present findings, together with these data, suggested that (1) the MET process in hair cells is not confined to the stereocilia but rather applies to other subcellular distributions, such as the apical surface, where the cells might perceive mechanical forces from the extracellular matrix or fluid motion; (2) the VHCs have different MET characteristics from the CHCs, which are at least partially attributed to specifically distributed mechanosensitive GPCRs. Although much remains unknown about the regulatory mechanism and physiological importance of the MET currents at the apical surface of VHCs, as well as their potential relationship with the MET currents at the stereocilia, the distinct MET processes at different subcellular distributions potentially imply greater complexity of equilibrioception than of auditory perception. A combined MET information from both stereocilia and apical surface may help to provide a more precise spatial resolution for positional sensation compared with hearing.

Reconstitution of LPHN2-TMC1 functional coupling in the heterologous system suggested that force sensation by LPHN2 could induce TMC1 activation through a conformational transition between these two membrane proteins. We observed coimmunostaining of LPHN2 and TMC1 at the apical surface of the utricular hair cells (Supplementary information, Fig. S9f). Notably, although the LPHN2-TMC1 pair is sufficient to convert extracellular stimuli into electrical signals in the heterologous system, we cannot exclude the potential involvement of other subunits or accessory proteins in vivo. For example, the conventional MET component LHFPL5 may be expressed at the apical surface of CHCs, while the chaperone receptor LGR6 used in the present study is endogenously expressed at the apical surface of VHCs^58^. Whether these molecules are part of the apical surface MET machinery requires further investigation. Moreover, since LPHN2 is expressed in approximately 80% of utricular hair cells, it is of interest to explore whether and how the MET process might be regulated by other mechanosensitive GPCRs in the remaining 20% of LPHN-negative hair cells. Future studies regarding these questions will provide in-depth insights into more functional roles of GPCRs in equilibrioception.

In addition to present in hair cells, the scRNA-seq datasets suggested that Lphn2 were expressed in the utricular supporting cells. However, using both a commercially available LPHN2 antibody and a transgenic knock-in mouse line, our results indicated that LPHN2, at the protein level, was specifically expressed in the utricular hair cells but was not detectable in the supporting cells. Currently, we don’t know the mechanism underlying the discrepancy of LPHN2 expression measured at the protein and at mRNA levels. We speculated that this difference might be due to different protein level controlling pathways, such as the different protein degradation system of LPHN2 proteins in supporting cells or hair cells, which need further investigation. Beyond the hearing and vestibular system, LPHN2 is widely expressed and regulates various functions, such as synapse formation^72^, heart development^73^ and vascular remodeling^74^. The diverse expression pattern and multiple function of LPHN2 is similar to the well-known force-sensitive ion channel Piezos. We speculate that the mechanosensitive channels or receptors might respond to diverse types of forces (e.g., cellular compression, fluid shear stress, membrane tension) in different organs or tissues. However, further studies are required to investigate the function of the force sensation by LPHN2 in other tissues or cells.

In conclusion, we identified a force-sensitive GPCR expressed at the apical surface of VHCs (1), which senses force within a physiological range (2), regulates a tip-link-independent MET process and converts force stimuli into neurotransmitter glutamate release (3). Specific ablation of this particular receptor in hair cells severely impairs the balance of mice (4). These results conform to the criteria we proposed for equilibrioception receptors and suggest an indispensable role of GPCRs in equilibrioception.

## Limitations of the Study

Our current study indicates that LPHN2 expressed at the apical surface of utricular hair cells regulates a tip link-independent MET current by converting force stimuli into TMC1 activity, which is indispensable for the normal equilibrioception. However, mechanisms underlying the different effects of BAPTA on fluid jet-stimulated MET in CHCs and VHCs are not known. Further studies using more precise tools to differentiate the tip link-dependent and -independent MET are required for the mechanistic investigation. In addition, although the magnetic-bead-delivered force in the present study is theoretically within the physiological range, its physiological significance was limited by the relatively low binding efficiency of the antibody-coated beads and the potential time delay in transmitting force. Further in-depth investigation of the functional roles of VHC-expressed LPHN2 in the equilibrioception requires the real-time recording of Ca^2+^ signaling and neurotransmitter release in VHCs in response to a natural force stimulus, such as fluid jet or stiff probe-induced deflection of hair bundles, using mouse models with Lphn2 deficiency or pharmacological blockade. Moreover, in vivo two-photon calcium or neurotransmitter imaging in moving mouse models is expected to further elucidate the equilibrioception potential of LPHN2 in a more physiological setting.

## Data availability

All data are available upon reasonable request from the corresponding authors.

## Acknowledgments

This work was supported by National Key R&D Program of China (2019YFA0904200 to J.-P.S., 2021YFA1101300 to R.-J.C., 2023YFA1801100 to Z.Y.), National Science Fund for Distinguished Young Scholars Grant (81825022 to J.-P.S., 82225011 to X.Y.), National Natural Science Foundation of China (T2321004 to J.-P.S. and F.Y., 32130055 and 82330118 to J.-P.S., 82330022 to X.Y., 82030029 to R.-J.C., 82271176 to W.-W.L.), the Key Research Project of the Beijing Natural Science Foundation, China (Z200019 to J.-P.S.), Major Fundamental Research Program of Natural Science Foundation of Shandong Province, China (ZR2020ZD39 to J.-P.S., Z.-G.X. and W.-W.L, ZR2021ZD18 to X.Y.), Taishan Scholars Program of Shandong Province (tsqn201909189 to W.-W.L.), Shandong Provincial Natural Science Fund (ZR2023QH394 to M.-W.W.) and the Cutting Edge Development Fund of Advanced Medical Research Institute, Shandong University (to J.-P.S., Z.Y. and K.-K.Z.). J.-P.S. is also supported by the Tencent New Cornerstone Investigator Program.

## Author contributions

J.-P.S., X.Y. and R.-J.C initiated, designed and supervised the overall project. J.-P.S., X.Y. and Z.Y. starting the screening of mechanosensitive GPCRs in vestibular and cochlear systems from 2019. R.-J.C, F.Y. and Z.-G.X. provided guidance and suggestions on the vestibular and cochlear studies. Z.Y. and J.-P.S. developed the magnetic force stimulation assay to examine the activation of G proteins and second messengers downstream of GPCRs. X.Y. designed and supervised all electrophysiology experiments. W.Y. provided guidance and suggestions on the electrophysiology experiments. Z.Y., S.-H.Z., Q.-Y.Z. and Z.-C.S. performed the cell-based assay of LPHNs. S.-H.Z., Z.Y., Q.-Y.Z. and Z.-C.S. performed the electrophysiological experiments and Ca^2+^ imaging supervised by J.-P.S. and X.Y.. J.-P.S., Z.Y., S.-H.Z. and W.-W.L. participated in the design of gene knockout mice and modified AAV-ie. Z.Y., S.-H.Z., Q.-Y.Z., Z.-C.S, and W.-W.L. performed vestibular behavior studies. Z.Y., S.-H.Z., Q.-Y.Z., Z.-C.S, Y.G., J.-Y.Q., X.-H.W. and Y.L. isolated mouse utricle and performed immunofluorescence studies and ex vivo experiments. Z.Y., M.-W.W., S.-H.Z., and Q.-Y.Z. performed single cell qRT-PCR. Y.G., M.-W.W., S.-H.Z. and Y.-N.S. performed coimmunoprecipitation and western blotting analyses. Z.Y., S.-H.Z., Q.-Y.Z. and W.-W.L. performed the AAV-ie-related experiments under the supervision of J.-P.S. and R.-J.C.. K.-K.Z. performed virtual screening under the supervision of J.-P.S.. J.-P.S., X.Y., R.-J.C., F.Y., W.Y., Z.Y., S.-H.Z., Q.-Y.Z., Z.-C.S. and Y.S. participated in data analysis and interpretation. W.Q. provided the Gauss/Tesla meter for the quantification of magnetic force. Y.S. provided insightful ideas related to the balance sensation based on clinical practice. Z.Y., S.-H.Z., Q.-Y.Z., Z.-C.S. and M.-W.W. prepared the figures. J.-P.S., X.Y., Z.Y. and R.-J.C. wrote the manuscript. All the authors have seen and commented on the manuscript.

## Competing interests

The authors declare no competing interests.

## Notes

### Competing Interest Statement

The authors have declared no competing interest.

### Summary of Updates

All figures updated Author list updated 1. A screening process for mechanosensitive GPCRs in the vestibular system has been supplemented. 2. Fluid-jet assay was employed to record the MET currents, which revealed an indispensable role of LPHN2 in normal MET response in utricular hair cells. 3. Electrophysiological studies were performed to support that the sensation of force by LPHN2 could enhance the activation of TMC1. 4. The author list has been changed according to their respective contributions to the revised version of the manuscript.

